# The relative contributions of infectious and mitotic spread to HTLV-1 persistence

**DOI:** 10.1101/799197

**Authors:** Daniel J Laydon, Vikram Sunkara, Lies Boelen, Charles R M Bangham, Becca Asquith

**Affiliations:** MRC Centre for Global Infectious Disease Analysis, Department of Infectious Disease Epidemiology, School of Public Health, Imperial College London, London W2 1PG, United Kingdom; Section of Immunology, Wright-Fleming Institute, Imperial College School of Medicine, London W2 1PG, United Kingdom; Department of Mathematics and Computer Science, Freie Universität, Arnimallee 6 14195 Berlin, Germany

## Abstract

Human T-lymphotropic virus type-1 (HTLV-1) persists within hosts via infectious spread (*de novo* infection) and mitotic spread (infected cell proliferation), creating a population structure of multiple clones (infected cell populations with identical genomic proviral integration sites). The relative contributions of infectious and mitotic spread to HTLV-1 persistence are unknown, and will determine the efficacy of different approaches to treatment.

The prevailing view is that infectious spread is negligible in HTLV-1 proviral load maintenance beyond early infection. However, in light of recent high-throughput data on the abundance of HTLV-1 clones, and recent estimates of HTLV-1 clonal diversity that are substantially higher than previously thought (typically between 10^4^ and 10^5^ HTLV-1^+^ T cell clones in the body of an asymptomatic carrier or patient with HAM/TSP), ongoing infectious spread during chronic infection remains possible.

We estimate the ratio of infectious to mitotic spread using a hybrid model of deterministic and stochastic processes, fitted to previously published HTLV-1 clonal diversity estimates. We investigate the robustness of our estimates using two alternative methods. We find that, contrary to previous belief, infectious spread persists during chronic infection, even after HTLV-1 proviral load has reached its set point, and we estimate that between 100 and 200 new HTLV-1 clones are created and killed every day. We find broad agreement between all three methods.

The risk of HTLV-1-associated malignancy and inflammatory disease is strongly correlated with proviral load, which in turn is correlated with the number of HTLV-1-infected clones, which are created by de novo infection. Our results therefore imply that suppression of de novo infection may reduce the risk of malignant transformation.

**Author Summary:** There are no effective antiretroviral treatments against Human T-lymphotropic virus type-1 (HTLV-1), which causes a range of inflammatory diseases and the aggressive malignancy Adult T-cell Leukaemia/Lymphoma (ATL) in approximately 10% of infected people. Within hosts the virus spreads via infectious spread (*de novo* infection) and mitotic spread (infected cell division). The relative contributions of each mechanism are unknown, and have major implications for drug development and clinical management of infection. We estimate the ratio of infectious to mitotic spread during the infection’s chronic phase using three methods. Each method indicates infectious spread at low but persistent levels after proviral load has reached set point, contrary to the prevailing view that infectious spread features in early infection only. Risk of disease in HTLV-1 infection is known to increase with proviral load, via mutations accrued from repeated infected cell division. Our analyses suggest that ongoing infectious spread may provide an additional mechanism whereby chronic infection becomes malignant. Further, because antiretroviral drugs against Human Immunodeficiency Virus (HIV) inhibit HTLV-1 infectious spread, they may reduce the risk of HTLV-1 malignancy.

## Introduction

Human T-lymphotropic virus type-1 (HTLV-1), also known as the human T cell leukaemia virus, infects an estimated 10 million people worldwide [1]. While the majority of infected individuals remain lifelong asymptomatic carriers (ACs), in ∼10% the virus causes either Adult T-cell Leukaemia/Lymphoma (ATL) [2] or a range of inflammatory diseases, notably a disease of the central nervous system called HTLV-1-associated myelopathy/tropical spastic paraparesis (HAM/TSP) [3]. HTLV-1 viral burden is quantified by the proviral load (PVL), defined as the number of HTLV-1 proviruses per 100 peripheral blood mononuclear cells (PBMCs). During the chronic phase of infection, PVL remains approximately constant [4, 5] within each host, but varies between hosts by over four orders of magnitude; a high PVL is associated with HAM/TSP [5, 6] and ATL [7].

HTLV-1 replicates in the host through two pathways: mitotic spread and infectious spread [8]. In mitotic spread, an infected cell divides to produce two identical “sister cells which carry the single-copy provirus integrated in the same genomic location as the parent cell. Infectious spread, or *de novo* infection, occurs when the virus infects a previously uninfected cell, and in this case the virus integrates in a new site in the target cell genome [Figure 1]. The combination of infectious and mitotic spread results in a large number of distinct clones of infected T-cells, each clone defined as a population of infected cells with a shared proviral integration site [9–11].

**Figure 1.**
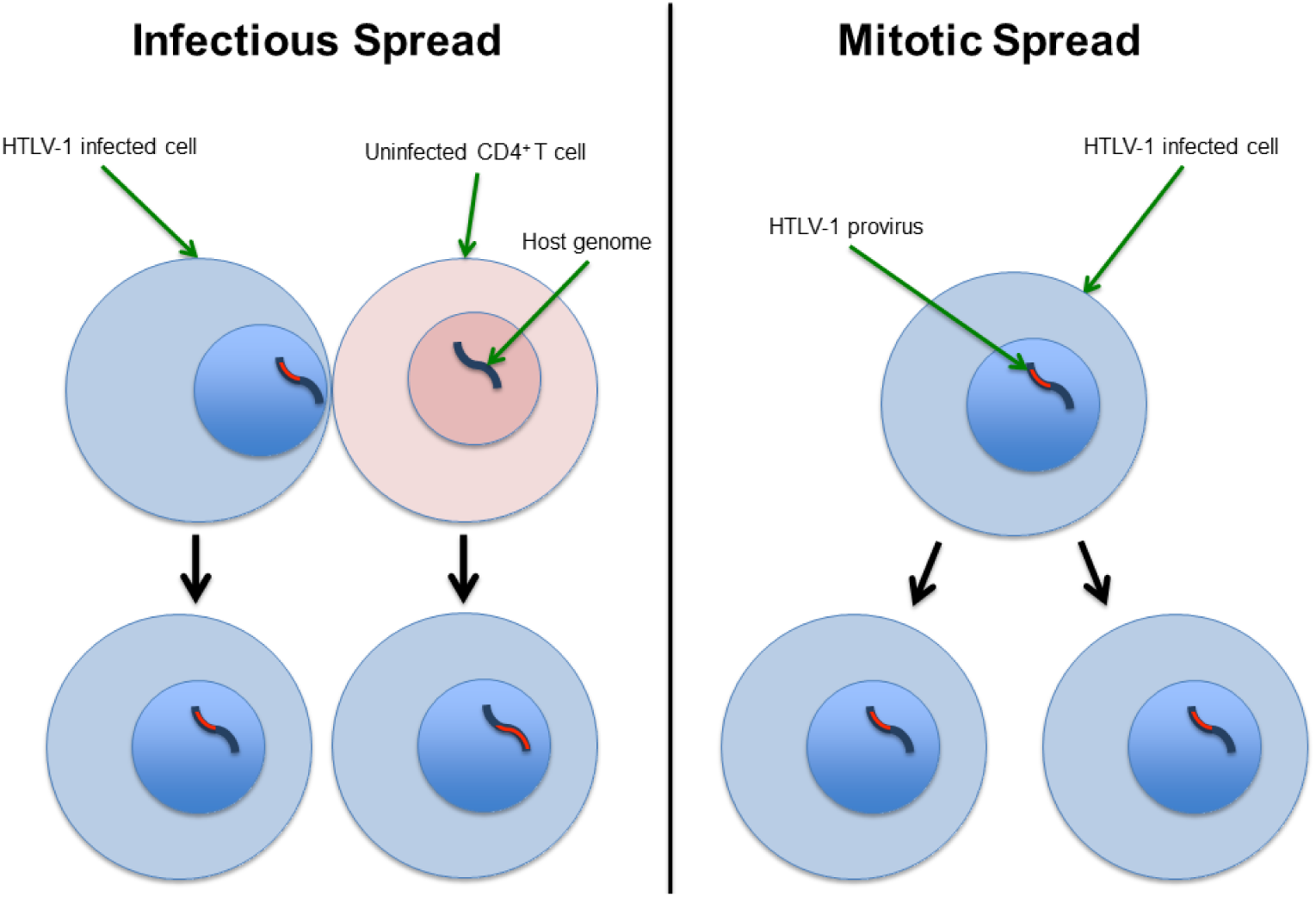
HTLV-1 infectious and mitotic spread schematic. Left column (Infectious spread): an HTLV-1-infected cell infects an uninfected CD4^+^ T cell (typically by cell-to-cell contact via the virological synapse, and potentially also via cell-free spread). The HTLV-1 provirus (red) integrates in a different genomic location in the newly infected cell, so infectious spread has resulted in two clones. Right column (Mitotic spread): An HTLV-1-infected cell divides, whereupon the provirus resides in the same genomic location in each daughter cell. The figure shows a single clone with two HTLV-1-infected cells.

The relative contribution of infectious spread and mitotic spread to the proviral load is unknown. This ratio is important, because it will directly determine the efficacy of different approaches to treatment. Although no effective antiretroviral drugs have yet been developed for HTLV-1 infection, antiretroviral therapy (ART), which efficiently reduces infectious spread in HIV-1 infection by inhibiting reverse transcription, viral maturation and proviral integration, may be effective in HTLV-1 infection if infectious spread contributes to the maintenance of HTLV-1 proviral load. Alternatively, immunosuppressive drugs such as ciclosporin which inhibit T cell proliferation would be expected to be more useful if mitotic spread [8] is the dominant mode of viral spread.

The number of clones of HTLV-1-infected T cells depends on the extent of infectious spread. In this paper, we refer to this number as the HTLV-1 clonal “diversity” (this term should not be confused with measures such as Shannon entropy or beta diversity). The diversity in one host is unknown, and estimating this number from blood samples is nontrivial. Diversity estimation is challenging given the nature of the HTLV-1 clone frequency distribution, where the majority of infected cells are contained in relatively few clones, and the majority of clones contain relatively few cells.

The prevailing view is that mitotic spread accounts for the majority of HTLV-1 persistence [11–14], and that infectious spread is negligible after initial infection [12, 13]. This belief is supported by three main observations. First, it was thought that there were relatively few (∼100) HTLV-1 clones in one host [9, 11, 13, 15–19]. Second, HTLV-1 varies little in sequence both within and between hosts [20]. Since the host DNA polymerase used in cell proliferation (mitotic spread) is much less error-prone than the viral reverse transcriptase used in infectious spread, a lack of sequence variation implies that infectious spread is rare. Third, many HTLV-1^+^ clones have been observed at multiple time points separated by several years [9, 17], and a long-lived clone is very unlikely to be maintained by repeated proviral integration through infectious spread at the same integration site, especially since there are no hotspots of HTLV-1 integration [9].

However, these three observations do not necessarily imply that infectious spread is negligible [14], particularly when we consider the total number of clones in the host and the very small proportion of clones that can be sampled. First, estimates of the number of clones have increased over time [9, 11, 13, 15, 17, 19], and current estimates give approximately 10^4^ - 10^5^ clones in the circulation of ACs and patients with HAM/TSP [10, 21, 22]. Second, apparent sequence uniformity may result from repeated detection of sister cells from a small number of expanded clones. That is, because of the limitations of sampling, there is a strong bias to detection of the large clones which expanded through mitosis. Finally, the repeated observation of specific clones over many years does not rule out persistent infectious spread. The observation of a temporary but dramatic PVL reduction in a patient with HAM/TSP following treatment with the reverse transcriptase inhibitor lamivudine [23] implies that infectious spread remains important in HTLV-1 persistence, at least in some cases.

Even when taking recent estimates of clonal diversity into account, there is still good reason to believe that mitotic spread is predominant, because the 10^4^ to 10^5^ clones (created by infectious spread) present in one host consist of approximately 10^11^ infected cells (maintained by mitotic spread). However, this consideration ignores the possibility that clones may be continuously created by infectious spread and killed by the immune response and natural death.

The aim of this study was to quantify the rate of infectious spread, and thus the ratio of infectious spread to mitotic spread during chronic infection. We first estimated HTLV-1 clonal diversity in 11 subjects using our previously developed method [10]. We next developed a deterministic and stochastic hybrid model of within-host HTLV-1 persistence that we fitted to clonal diversity estimates. We further used two alternative approaches to quantify the rate and to ensure robustness of our estimates. First, we developed a simplified model to approximate the upper bound of the rate. Second, we adapted a method originally developed to model naïve T cell dynamics. We find broad agreement between estimates from all methods. We conclude that, during chronic infection, a given HTLV-1-infected cell in the peripheral blood is substantially more likely to be derived by mitosis of an existing clone than by de novo infection, although infectious spread continues throughout chronic infection with an average of 175 new clones created every day.

## Methods

### Data sets

We apply all three methods described below to previously obtained high-throughput data on HTLV-1 clonality [9]. Each HTLV-1 dataset quantifies the abundance of HTLV-1-infected T cell clones in ex vivo peripheral blood mononuclear cells, without selection or culture. We studied 11 subjects, where each subject had three blood samples taken per time point, at three time points separated by an average of 4 years, giving a total of 99 datasets. All subjects either had HAM/TSP or were asymptomatic carriers of HTLV-1.

### HTLV-1 clonal diversity estimates

To estimate the rate of infectious spread we first estimated HTLV-1 clonal diversity. We use our recently developed estimator, “DivE” [10, 24, 25], which uses experimental measurements of clonal diversity in a sample to estimate both the number of clones and their frequency distribution in the body of the host [Figure 2A]. DivE fits multiple mathematical models to individual-based rarefaction curves; such curves plot the expected number of clones against the number of infected cells sampled. Numerical criteria score models on their ability to accurately estimate additional data. The best-performing models are extrapolated to estimate the total number of clones in the body, based on the proviral load in each respective subject. See [10, 25] for further details and implementation.

**Figure 2:**
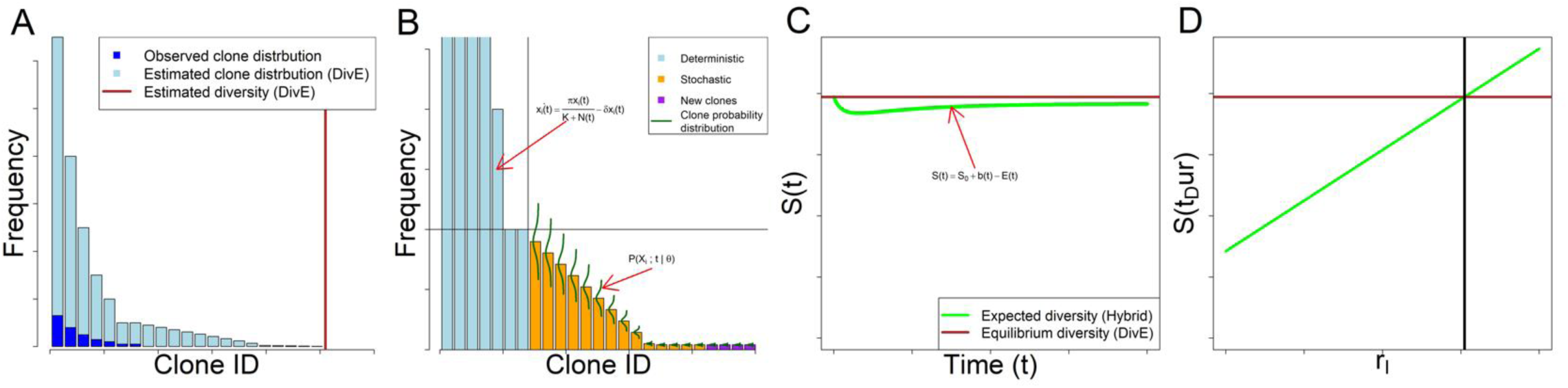
Schematic of full simulation hybrid model. **A**: Observed and estimated clone frequency distributions. From an observed sample of clones, the clone frequency distribution of the body in one host is estimated using DivE. **B**: Propagation of hybrid model: Estimated clone frequency distribution partitioned into deterministic and stochastic systems. Clones of frequency less than and greater than threshold *F* are respectively modelled stochastically and deterministically. *F* is chosen with respect to probability of clone extinction [supplementary information]. The deterministic system is modelled using ordinary differential equations [Eq. (1)]. The stochastic system consists of multiple birth-death processes (one for each stochastically modelled clone) each with an absorbing state at zero [Figure 3]. The evolution of the clone probability distribution over time is governed by the chemical master equation [Eq. (10), Figure 4]. New clones are created through infectious spread, i.e. the per-capita rate *r*_*I*_ multiplied by the expected number of infected cells, in both deterministic and stochastic compartments [Eq. (11)]. Deterministic and stochastic systems are propagated concurrently with Strang splitting [supplementary information]. **C**: Hybrid model diversity. The estimated number of clones *S(t)* [Eq. (13)] at time *t*, given parameters *θ = {π, δ, K, rI}* is given by the number of clones created [Eq. (11)], minus the number of clones that are expected to have died between 0 and *t* [Eq. (12)], plus the number of clones *S*_*0*_ at *t = 0*. The number of clones is assumed to be at equilibrium in the chronic phase of infection. **D**: Model fitting schematic: Expected diversity at *S(t*_*Dur*_) increases with per-capita infectious spread rate *r*_*I*_. Model fitted using non-linear least squares to DivE estimated diversity in the body, where the objective function is the square of the discrepancy between this value and the value of *S(t*_*Dur*_) at equilibrium.

Table S1 gives the notation used in the three modelling approaches that follow.

### Modelling approach 1: Full simulation hybrid model

Within a given host, HTLV-1^+^ T cell clones vary in abundance by several orders of magnitude [9, 10]. Broadly, abundant clones can be modelled deterministically but small clones must be modelled stochastically. In the following sections, we describe a model of HTLV-1 dynamics at quasi-equilibrium that is a hybrid of deterministic and stochastic parts [Figure 2].

#### Deterministic Model

We consider a system with *S(t)* clones, where a given clone *i* has frequency *x*_*i*_*(t)* at time *t*. We have the following ordinary differential equations (ODEs) for each clone:

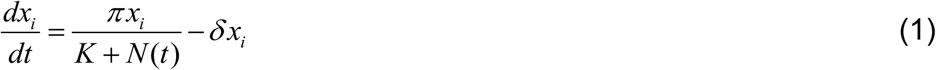

where 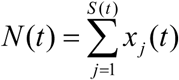 is the total number of infected cells summed over all clones at time *t*; 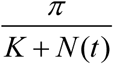 is the proliferation rate of infected cells (i.e. the rate of mitotic spread) which is half maximal when *N(t) = K* (see supplementary information) and *δ* is the death rate of infected cells [Figure 2B].

The dynamics of small clones, where random effects are important, will not be adequately described by a deterministic model. Since small clones contain most information about infectious spread, it is important to model these clones accurately, and so we use a discrete stochastic model, in which we consider multiple potential states of each clone and their corresponding probabilities over time.

#### Stochastic Model

Using a stochastic framework, the number of clones *S(t)* and their frequencies at time *t* are considered as random variables, and we describe within-host HTLV-1 dynamics by a set of reactions and their corresponding propensities [supplementary information]. Infected cells can proliferate, die, or infect uninfected cells [Figure 1]. Thus the total number of possible reactions *C* ∈ ℕ at time *t* is *C = 3S(t)*. Following the formulation given in [26, 27], let *X* (*t*) = ((*X*_*i*_ (*t*)) _*i*∈*S*(*t*)_)^*T*^ be the state vector at time *t* of all clones. *X(t)* is a random variable in 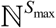 that consists of the random variables *X*_*i*_ (*t*) ∈ ℕ_0_ = ℕ ⋃{0} of the frequencies *x*_*i*_*(t)* of clones *i =* 1,…, *S*_*max*_, where *S*_*max*_ is chosen to always be larger than *S(t)* for all *t*. The state vector *X(t)* evolves through a Markov jump process that depends only on the current state 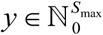, and its evolution is given by

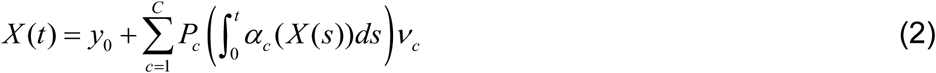

where *ν*_*c*_ and *α*_*c*_ respectively denote the stoichiometric vector and propensity function of reaction *c* [26, 27]. Equation (2) states that the population *X(t)* at time *t* is equal to the initial population *y*_*0*_ plus the sum of the changes induced by all reactions. See supplementary information for further details.

There exists a probability distribution associated with the random variable 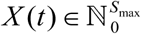 in (2), given by ℙ (*X*;*t*) = ℙ (*X* (*t*) = *y* | *X* (0) = *y*_0_), where 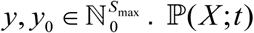 is a column vector where each entry is a probability associated with a potential state of the random variable at time *t*. It can be shown [27–30] that ℙ(*X*;*t*) is a solution of the Chemical Master Equation (CME)

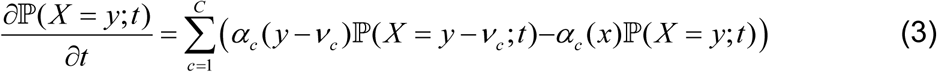

which describes the rate of change in the probability distribution associated with *X(t)*. The first term is the sum over all reactions of the probability of arriving at state *X(t) = y* from state *X(t) = y - V*_*c*_ via reaction *c*, and the second term is the sum over all reactions of the probability of leaving state *X(t) = y* via reaction *c*.

For a single clone *𝒳*_*i*_, the following reactions respectively describe mitotic spread, cell death and infectious spread:

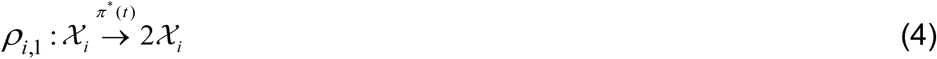

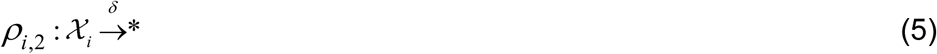

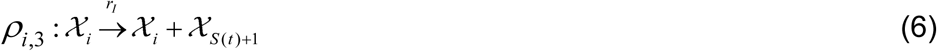

where

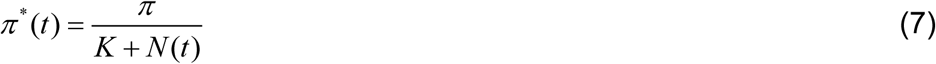

is the aggregate density-dependent proliferation rate (dependent on the carrying capacity, and the numbers of infected and uninfected cells). The first two reactions of each clone describe a birth-death process, and the lack of inflow from source (i.e. the lack of a reaction *ρ*:* →*𝒳*_*i*_) defines an absorbing state [Figure 3].

**Figure 3.**
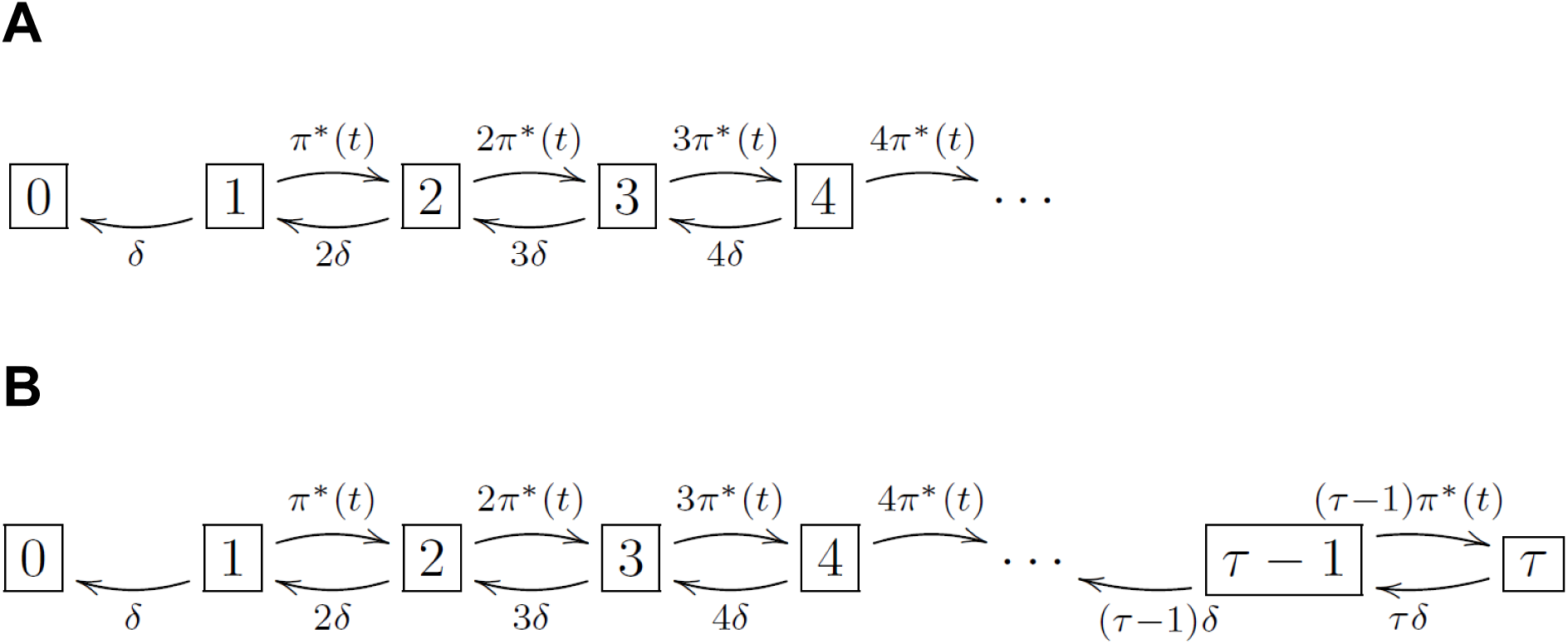
Clone state space birth-death process flow diagram. Each box denotes the potential state of a given clone, i.e. the number of cells in that clone, with the corresponding propensity of each reaction at each state. *π*\**(t)* and *δ* denote the per-capita rates of infected cell proliferation and death respectively. Note there is no source inflow from frequency *0* to frequency *1*. **A** and **B** respectively show the state space with and without an upper limit *τ.*

The reactions (4), (5) and (6) are monomolecular (in terms of the chemical master equation), because they carry the simplifying assumption that cell death due to the host immune response, and the proviral load, are each constant in the equilibrium within each host. HTLV-1 proviral load remains stable over many years [4, 5]: that is, the numbers of infected and uninfected cells stays approximately constant during the chronic phase of infection.

#### Simplifying approximations of stochastic model

The probability distribution ℙ(*X*;*t*) describes the states and associated probabilities of the entire system, and we define the probability distribution of a particular clone *I* ℙ(*X*_*i*_;*t*) associated with the random variable *X*_*i*_*(t)* similarly: ℙ(*X*_*i*_; *t*) = ℙ(*X*_*i*_ (*t*) = *x*_*i*_ | *X*_*i*_ (0) = *x*_*i*,0_), where *x*_*i*_, *x*_*i*,0_ ∈ ℕ_0_. The extinction probability of clone *i* at time *t*, ℙ(*X*_*i*_ = 0;*t*), will be used below to calculate the expected number of clones at time *t* [Figure 2C], which in turn will enable our model to be fitted to HTLV-1 clonal diversity estimates [Figure 2D].

If clones interact and are modelled with a single master equation associated with ℙ(*X*;*t*), the complexity and runtime of the model increase exponentially with the number of clones. However, because we model the system when proviral load is in equilibrium and can therefore use monomolecular reactions, density-dependent proliferation rates remain approximately constant, and so we can model each clone in isolation with multiple master equations associated with multiple clone-specific distributions ℙ(*X*_*i*_;*t*) (*i =* 1, …, *S(t)*) [Figure 2B]. Therefore, the model complexity and runtime increase only linearly with the number of clones.

If we impose a maximum frequency for a particular clone *i* (supplementary information) [Figure 3B], we can summarise Equation (3) using multiple, simpler differential equations below

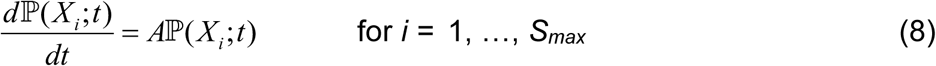

where *A* is the transition matrix or “matrix of connections” [supplementary information] [27, 31, 32]. Further, because the proliferation rate is constant at equilibrium, rates are independent of time, and so Equation (8) has solution

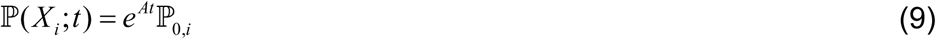

where ℙ_0,*i*_ = ℙ(*X*_*i*_;*t* = 0) is the initial probability distribution and *e*^*At*^ is the matrix exponential [33]. For equally spaced time steps 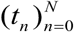 of length *h*, ℙ(*X*_*i*_;*t*) can be calculated recursively

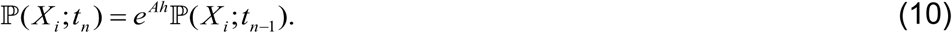

Example solutions of Equation (9) are shown in Figure 4.

**Figure 4.**
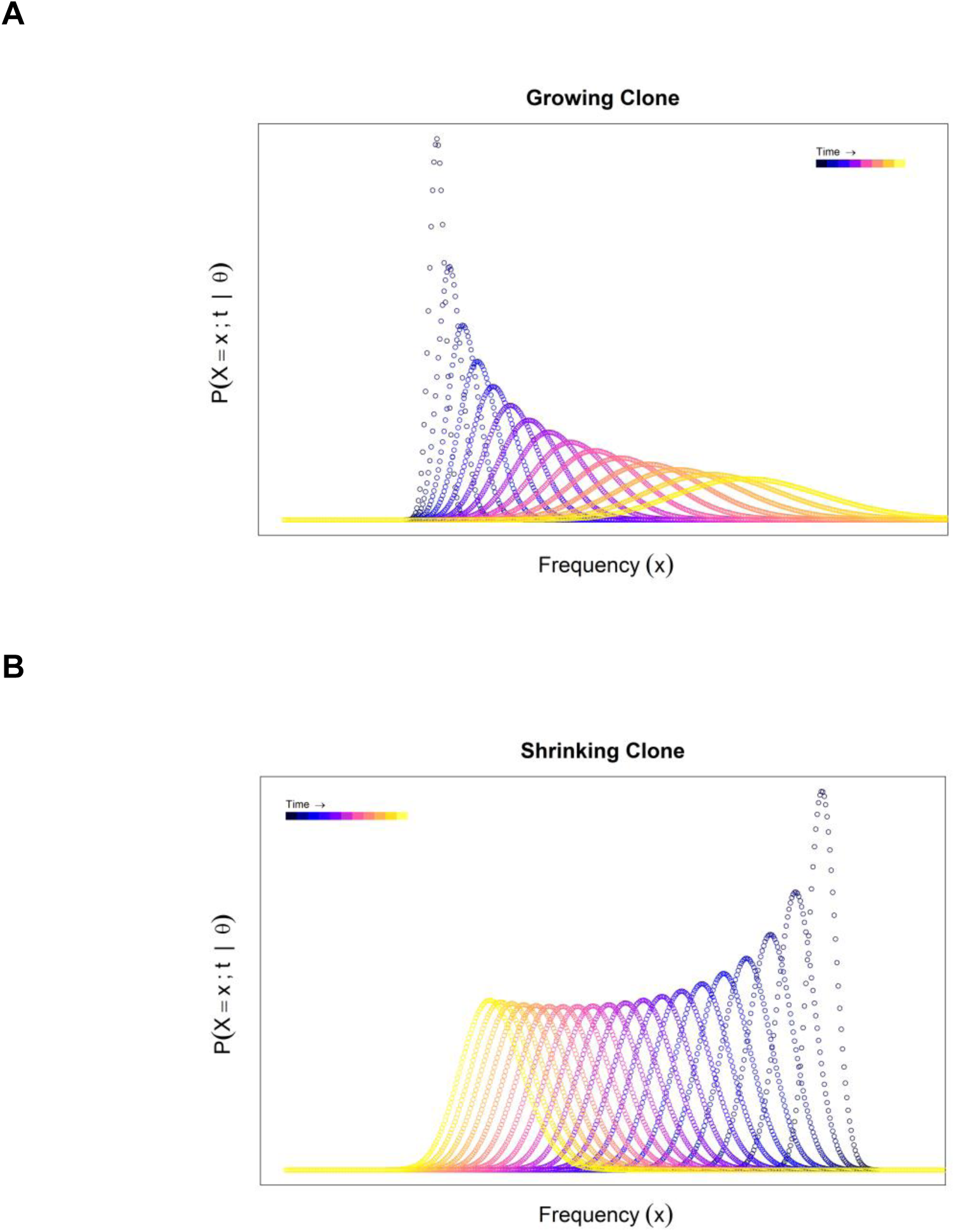
Probability distribution evolution. Each curve shows the distribution ℙ (*X*_*i*_; *t*) = ℙ (*X*_*i*_ (*t*) = *x*_*i*_ | *X*_*i*_ (0) = *x*_*i*,0_) of the probability that the given clone *i* contains *x*_*i*_ cells at time t. At successive time points the curve broadens and either **(A)** shifts to the right as the expected frequency of the clone increases, or (**B)** shifts to the left as the expected frequency of the clone decreases.

#### Expected number of clones

We model the expected number of clones *S(t)* at time *t* using by adding the total number of clone “births” *b(t)* over time (that is, the number of infectious spread events), and subtracting the total number of clone extinctions *E(t)* over time. *b(t)* is given by

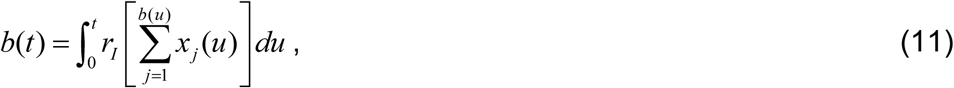

where *r*_*I*_ is the per-capita rate of infectious spread, *x*_*j*_*(t)* is the expected frequency of the *j*^th^ clone to be born since *t* = 0 (i.e. *x*_*j*_ (*t*) =𝔼[*X* _*j*_ (*t*)]), and *b(*0) = 0. *E(t)* is then given by

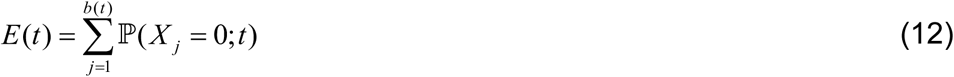

Note that *b(t)* and *E(t)* are increasing functions since *r*_*I*_, *x*_*j*_*(t)* ≥ 0, and because a clone frequency of zero is an absorption state for the random variable *X*_*j*_*(t)*. Taking (11) and (12) together we calculate the number of clones *S(t)* as

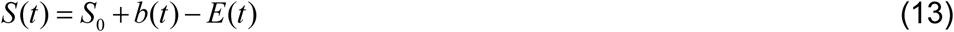

where *S*_*0*_ is the number of clones at time zero [Figure 2C].

#### Hybrid model fitting and uncertainty

It is estimated that there are approximately 10^11^ HTLV-1 infected cells in one host [10], and so it is not computationally feasible to model all clones using our stochastic formulation. Clones above a certain frequency [*F* = 460 cells; supplementary information] are assumed to be adequately described by the expected value from the deterministic ODEs in Eq. (1) [Figure 2B-D]. We thus partition our system of HTLV-1 within-host dynamics into a deterministic system of ODEs, and a stochastic system of master equations [Figure 2B]. We propagate these systems alternatively and concurrently using “Strang splitting” [supplementary information] [34]. The deterministic system described in Equation (1) has *S(t)* ordinary differential equations. Since the *S(t)* can exceed 10^5^, we group clones into categories based on the order of magnitude of their abundance.

We model the dynamics of clones in the body, and not only the blood, because this allows us to model clone extinction. If zero cells of a particular clone are observed or estimated in the blood, this does not necessarily imply that the clone is extinct, because cells in that clone could remain in the solid lymphoid tissue, which contains 98% of lymphocytes. We model clones in the body as a whole to avoid this difficulty, which necessitates the assumption that the clonal population structure in the blood is representative of the HTLV-1 clonal structure in the whole body.

We fitted the infectious spread rate *r*_*I*_ as a free parameter, with all other parameters (infected cell proliferation rate, death rate and density dependency) fixed using previous results from the literature and based on each subject’s proviral load [35] [supplementary information]. For each subject sample and parameter update of *r*_*I*_, the model was run to reach an approximate equilibrium [Figure 2C]. The model was fitted to the estimated clonal diversity of that subject sample, i.e. to determine the value of *r*_*I*_ required to keep the clonal diversity at the observed equilibrium value [Figure 2D].

The uncertainty in the estimate of *r*_*I*_, the rate of infectious spread, derives from three sources: error in model choice (both structure and numerical value of fixed parameters), error in clonal diversity estimation, and sampling variation. Classical methods of quantifying fitted parameter uncertainty only reflect the last source of error (i.e. they assume that the model and the data are correct). We address the first difficulty by using three alternative models with different structures and parameters. We address the error in diversity estimation by using alternative clonal diversity inputs from the Chao1 estimator [36], a non-parametric diversity (or species richness) estimator that has been widely used in many fields [37–40]. And we address the issue of sampling variation by investigating the range of estimates provided by the nine hybrid model fits per subject (i.e. one for each of the subject’s blood samples); the mean of these estimates is taken as our point estimate.

The hybrid model was coded in R (version 3.5.0) [41], using the packages “data.table [42] and “Matrix” [43]. Matrix exponentials were computed using the Padé approximation [44]. The hybrid was fitted using one-dimensional optimisation as described in [45].

### Modelling approach 2: upper bound approximation

We considered a simplified model of HTLV-1 persistence that does not describe individual clone dynamics. If *S(t)* and *N(t)* are the number of clones and number of infected cells respectively at time *t*, and *r*_*I*_, is the per-capita rate of infectious spread, we have the following differential equation

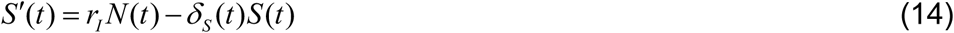

where *δ*_*S*_*(t)* is the *clone* death rate at time *t.* The first term of Equation (14) models the birth of new clones by infectious spread, and the second term models the death of existing clones.

If *δ* is the (constant) death rate of infected *cells*, then we have *δ*_*S*_*(t)* ≤ *δ*, because the number of clones that die cannot exceed the number of cells that die (equality would occur if all clones were singletons i.e. clones that contain only one infected cell). The clone death rate depends on the population structure of infected cells and will vary over time as this population structure changes. For example, a higher proportion of singletons will increase *δ*_*S*_*(t)*.

We assume that, in the chronic stage of infection when HTLV-1 proviral load is at equilibrium, the number of clones is also at equilibrium and so we have *N(t) = N, S’(t) = 0*, and *S(t) = S*. Letting *δ*_*S*_ be the average rate of clone death, we can approximate Equation (14) as

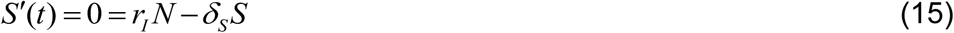

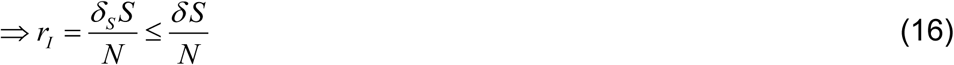

and therefore we define the supremum of the rate

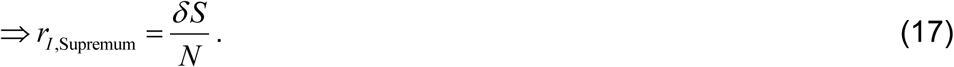

*r*_*I,Supremum*_ will substantially overestimate infectious spread because it applies the relatively high singleton death rate to all clones (clones with few cells become extinct more quickly than clones with many cells). To obtain a tighter upper bound we divide clones into those smaller and larger than an arbitrary size *f*_*max*_ and expand the expression for *r*_*I*_ in Equation (17) to obtain

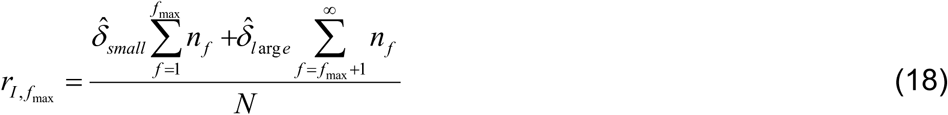

where *n*_*f*_ denotes the number of clones of frequency *f*, i.e. the “occupancy classes”. The aggregate clone death rate of small clones 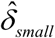 and of large clones 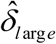 will comprise a weighted average of the death rate of clones of all sizes within that category. Because the HTLV-1 clonal frequency distribution is heavy tailed, small clones are more numerous than large clones, and so will make the dominant contribution to the clone death rate. Therefore the contribution from large clones can be neglected to give

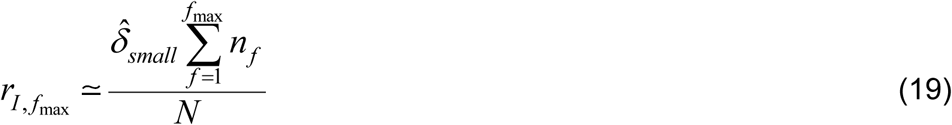

Provided *f*_*max*_ is sufficiently small, then 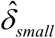 (which is less than or equal to δ) can be approximated by δ. The error incurred by this approximation decreases as *f*_*max*_ is reduced, and so the infectious spread rate will be best approximated by 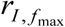 for low values of *f*_max_. Estimates of the ratio of infectious spread to mitotic spread can be obtained by dividing *r*_*I,Supremum*_ and 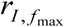 by the per-capita rate of mitotic spread *π =* 0.0316 [supplementary information] to give

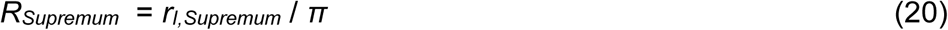

and

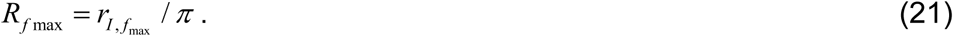

### Modelling approach 3: Occupancy class model

Adapting an model of naïve T cell dynamics [46], we model the occupancy classes *n*_*f*_ of HTLV-1 clones [Figure 5]. We assume that the clonal structure is in equilibrium (i.e. that the number of clones in each size class is constant) and that the probabilities of cell proliferation and death are independent of clone size.

**Figure 5.**
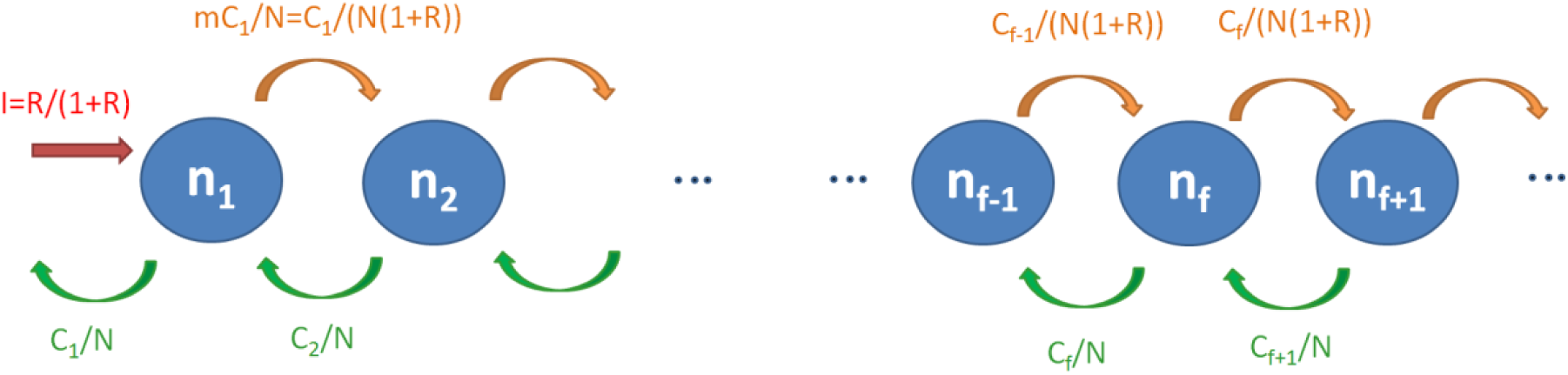
Occupancy class model schematic. Singletons (clones of size 1) are produced by infectious spread (red). Proliferation (orange) results in loss from clone size class *f* and entry into size class *f +* 1. Death of a cell (green) results in a clone moving from size class f to size class *f* - 1.

Scaling so there is one event (i.e. de novo infection or mitosis) per cell per unit time we have *I* + *M = 1* and *R := I / M.* Therefore

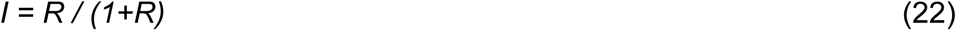

and

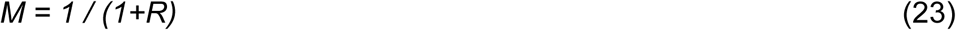

where *I* and *M* are the rates of infectious and mitotic spread (scaled as above), and *R* is the ratio of infectious to mitotic spread.

A clone in occupancy class *f* moves to class *f*+1 by mitosis with probability

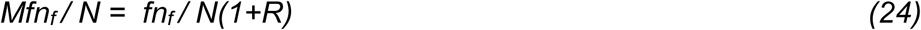

where *N* is the number of infected cells. A clone in occupancy class *f*+1 moves down to class *f* by death. Loss of cells by death is equal to the production of new cells by infection and mitosis, which has been scaled to 1, so the death rate is 1 per unit time. Since we assume that the probability of death is independent of clone size, the probability that the one death event in unit time occurs to a cell in size class *i*+*1* is simply equal to the proportion of cells in size class *i*+*1* i.e. *fn*_*f*_ */ N*.

In order for the number of cells *C*_*f*_ in size class *f* (*C*_*f*_ = *fn*_*f*_) to remain constant we require that flow in and flow out of the occupancy class *n*_*f*_ to be equal [Figure 5], i.e. that the number of cells leaving occupancy class *n*_*f*_ must be equal to those arriving from class *n*_*f-1*_ (via mitosis) and class *n*_*f*+*1*_ (via cell death). We therefore have

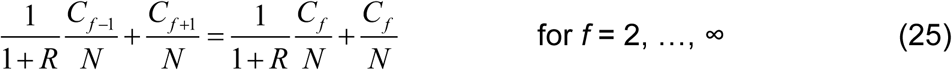

Rearranging gives

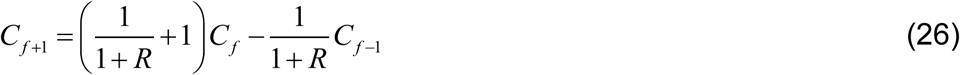

For the number of cells (*C*_*1*_) in size class 1 to remain constant we require

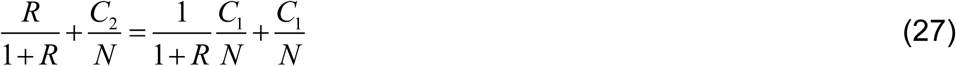

And for the population as a whole to remain of constant size we need the gain of new clones to balance their loss

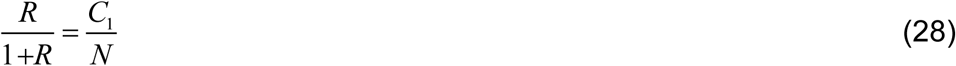

Rearranging (28) gives our first estimator (*R*_*1*_) for the ratio *R* from the occupancy class model, given in terms of *p =C*_*1*_*/N*, the proportion of cells that are singletons:

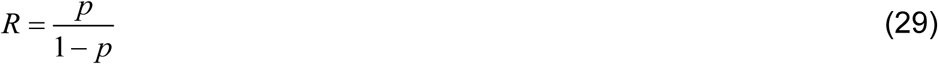

Substituting (28) into (27) and applying (26) recursively we obtain

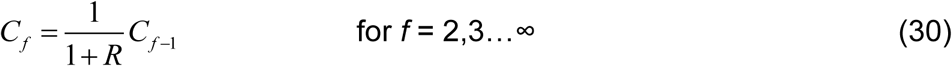

and thus

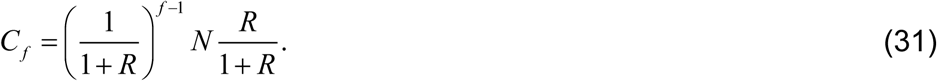

Species richness is defined as the number of clones, and so

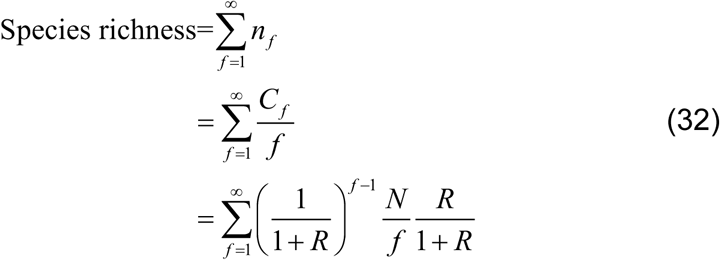

obtained by substituting in (31).

Using the fact that 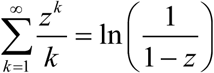 (a special case of the polylogarithm function)

We have that

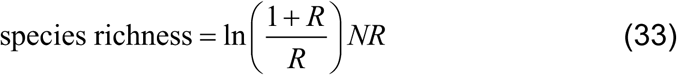

This is our second estimator for the ratio of infectious to mitotic spread, *R*_*2*_, from the occupancy class model.

The proportion of infected cells that are singletons is estimated using DivE, and the number of infected cells in the body is estimated from each patients proviral load as described in [10].

## Results

### HTLV-1 clonal diversity estimates

We estimated HTLV-1 clonal diversity (the number of unique clones) in 11 subjects with non-malignant HTLV-1 infection, either asymptomatic carriers or those with HAM/TSP. These estimates were obtained by measuring diversity in the nine blood samples per person (three at each of three time points) and then applying our recently developed method of estimating clonal diversity by extrapolation from the sample to the whole body [10] [Table 1].

**Table 1.**
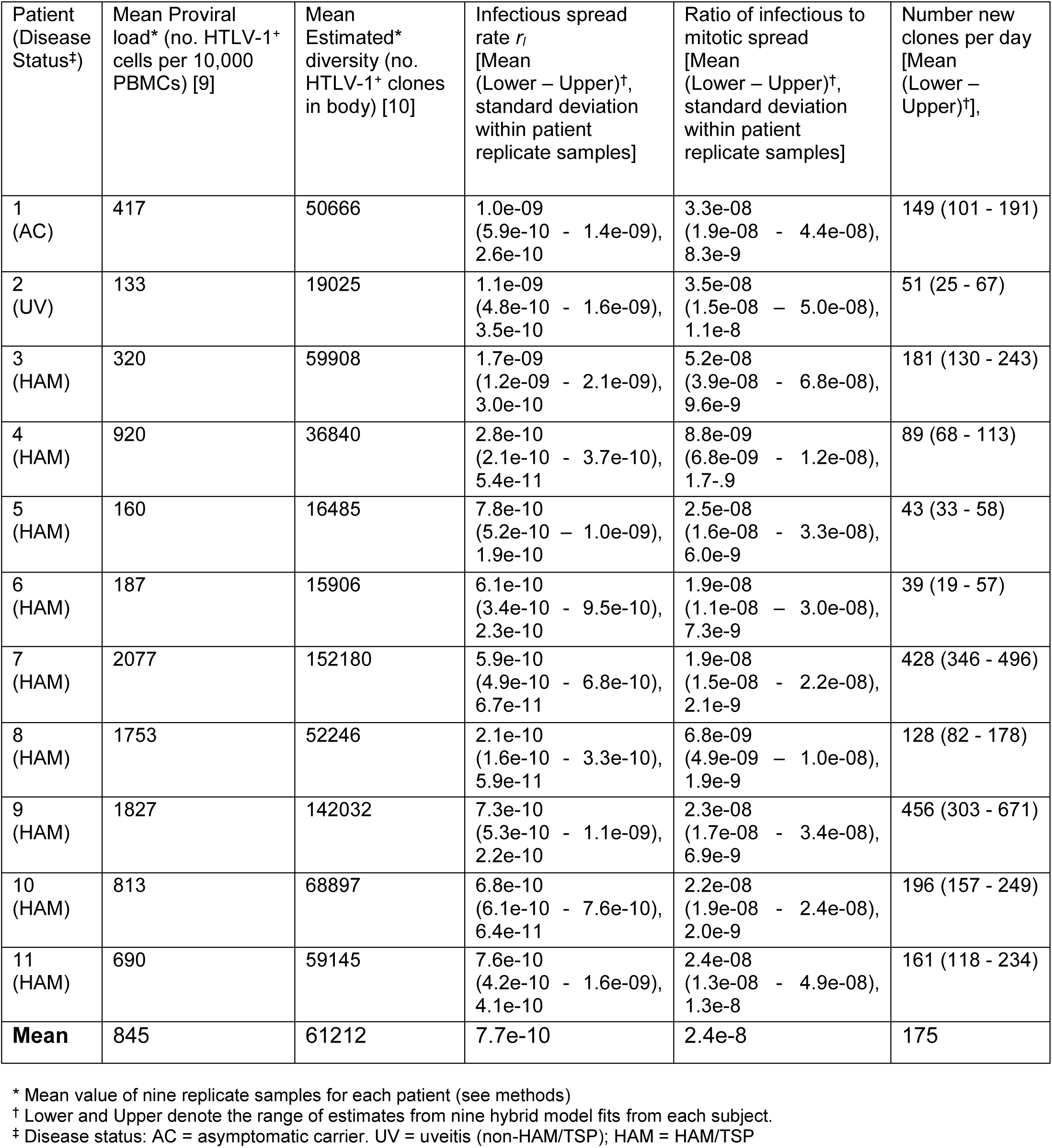
Hybrid model estimates of rate of infectious spread estimates and ratio of infectious to mitotic spread by patient.

**Table 2.**
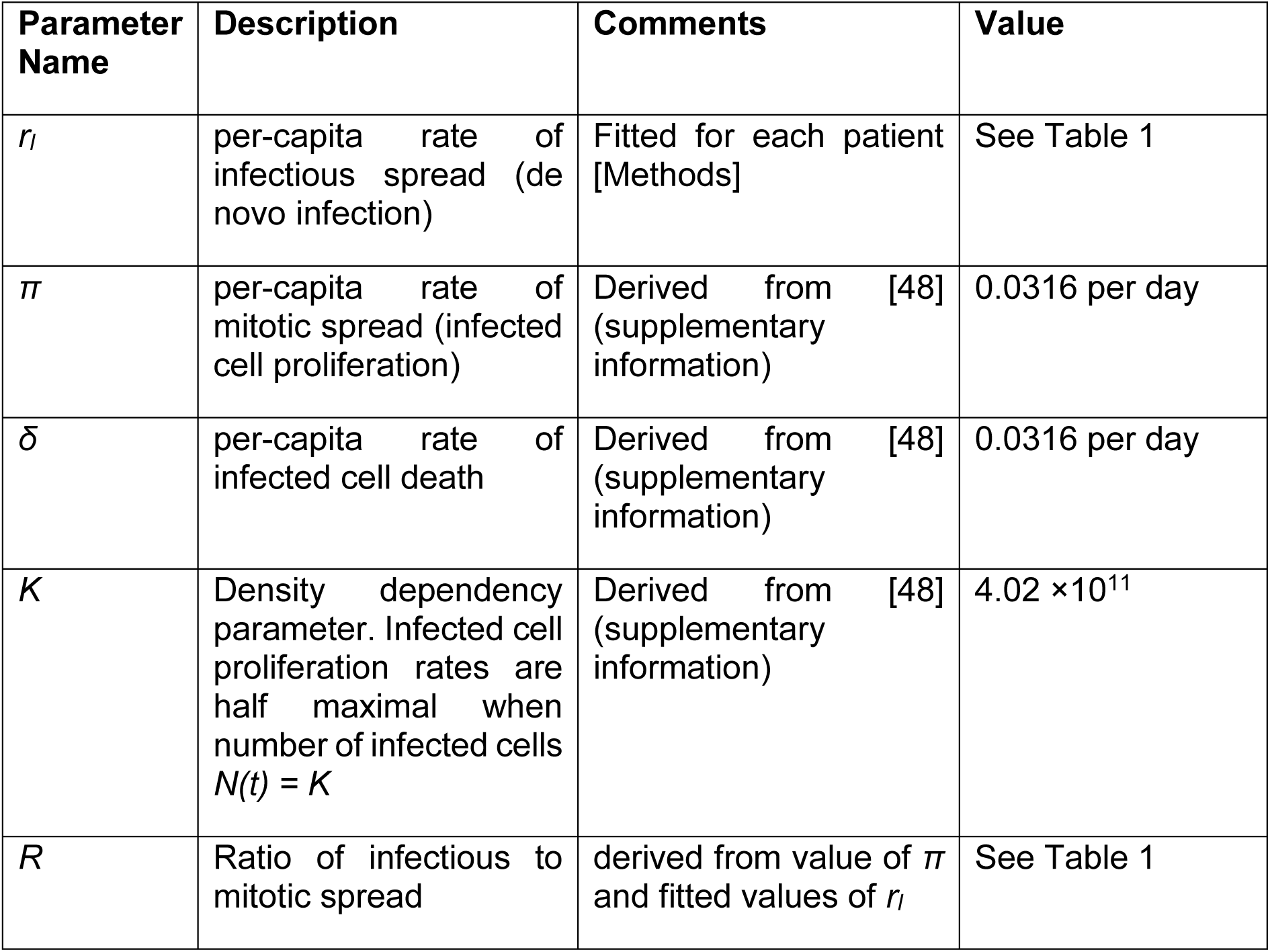
Parameter names and values

We tested our assumption that the number of clones is at equilibrium in the chronic phase of infection, where HTLV-1 proviral load is at equilibrium. We used linear regression to estimate the net change per day in the observed and estimated number of clones. This net change was 0.01 (95% CI -0.07 – 0.09) clones per day (i.e. 1 clone every 100 days) and -2.50 (−5.94 – 0.93) clones per day in the observed and estimated number of clones respectively; in each case the confidence interval spans zero. Further, using a two-tailed binomial test, we found little evidence that this change was significantly different from zero (p = 1 for observed and p = 0.07 for estimated). We therefore make the approximation that HTLV-1 clonal diversity remains unchanged in the chronic phase of infection, after the proviral load has reached steady state.

### Modelling approach 1: Full simulation hybrid model

Within-host HTLV-1 persistence is modelled by considering HTLV-1-infected clones individually. Large clones are modelled deterministically using a system of ordinary differential equations, whereas smaller clones are modelled stochastically by solving the chemical master equation [Equations (9) and (10)] that considers the frequency of each clone as a random variable governed by a birth-death process [Figure 2B]. The per-capita rate of infectious spread and the expected number of infected cells are then combined to model the birth of new clones (11), whereas the extinction probability of each clone is used to calculate expected clone death (12). The birth and death (or extinction) of clones provide an estimate of the number of clones at equilibrium (13) [Figure 2C], and it is this value that is fitted to our estimates of HTLV-1 clonal diversity, to infer the per-capita rate of infectious spread [Figure 2D].

The hybrid model was fitted to clonal diversity estimates for each subject (for each sample and each time point), providing an estimate of the infectious spread rate in each case [Table 1]. These nine estimates per patient were averaged to calculate the mean rate for each individual. Between individuals, the mean estimated rate of infectious spread was 7.7 × 10^−10^ per day, ranging from 2.1 × 10^−10^ to 1.7 × 10^−9^ per day [Figure 6A], i.e. varying by almost an order of magnitude. While this per-capita rate is very low, it translates to an average of 175 (range 39 - 456) new clones created per day [Figure 6B]. Therefore the hybrid model predicts that infectious spread is not limited to initial infection, but persists at a low level throughout the chronic phase. Given an estimate of the rate of mitotic spread of 3.2 × 10^−2^ per day, our infectious spread estimates imply an average ratio of infectious to mitotic spread of 2.4 × 10^−8^ (6.6 × 10^−9^ – 5.3 × 10^−8^) [Figure 7].

**Figure 6.**
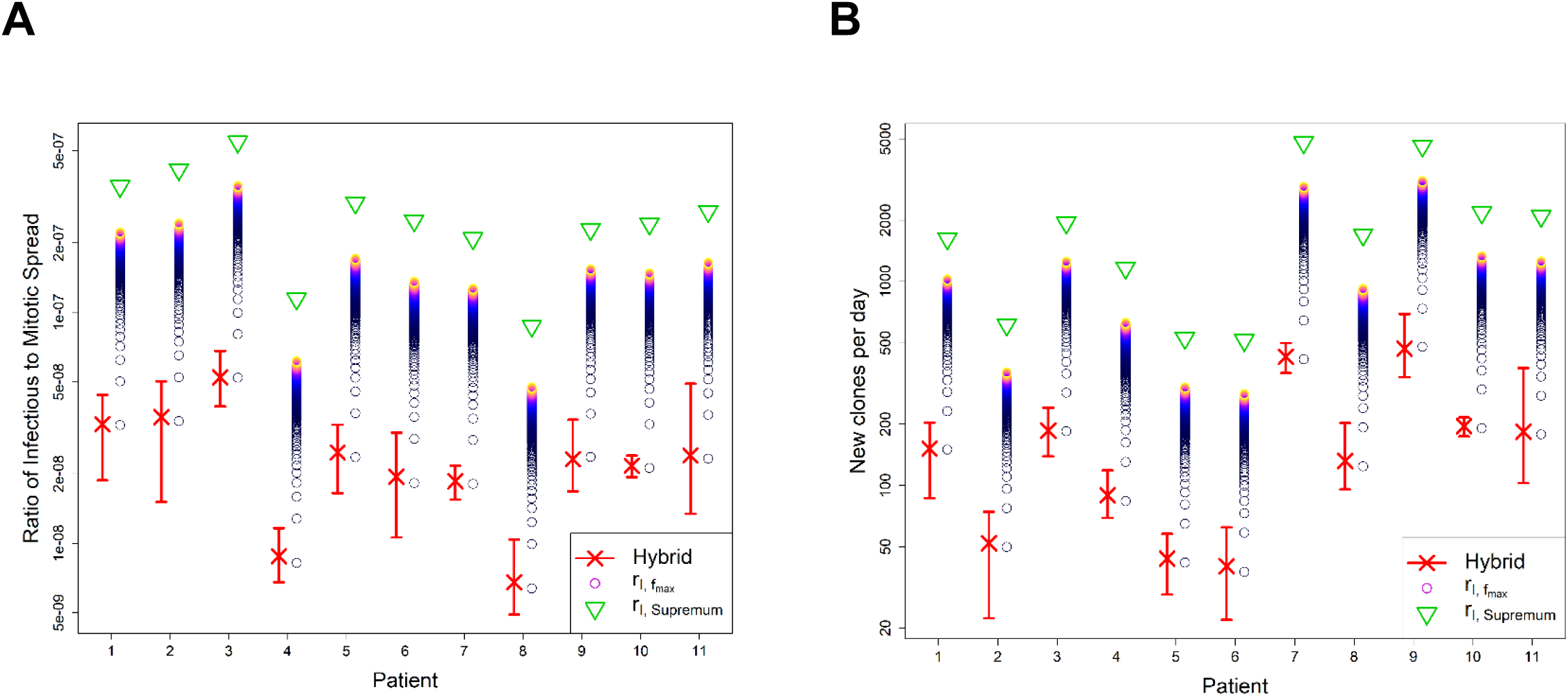
Ratio of infectious spread to mitotic spread and number of new clones per day, by patient and estimator. **A** Ratio of infectious spread to mitotic spread. **B** Number of new clones generated per day. In each plot, red crosses and bars respectively denote point estimates and the range from the nine estimates for each subject from the hybrid model. Upper bound approximations from *r*_*I,Supremum*_ (green triangles) are shown, together with tighter upper bounds from 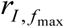 (coloured circles) for multiple values of *f*_max_ between 1 and 1000. Lighter colours denote higher values of *f*_max_. Hybrid model point estimates are very close to the estimates obtained for *f*_max_ = 1 (lowest circles). Estimates plotted on logarithmic scale.

**Figure 7.**
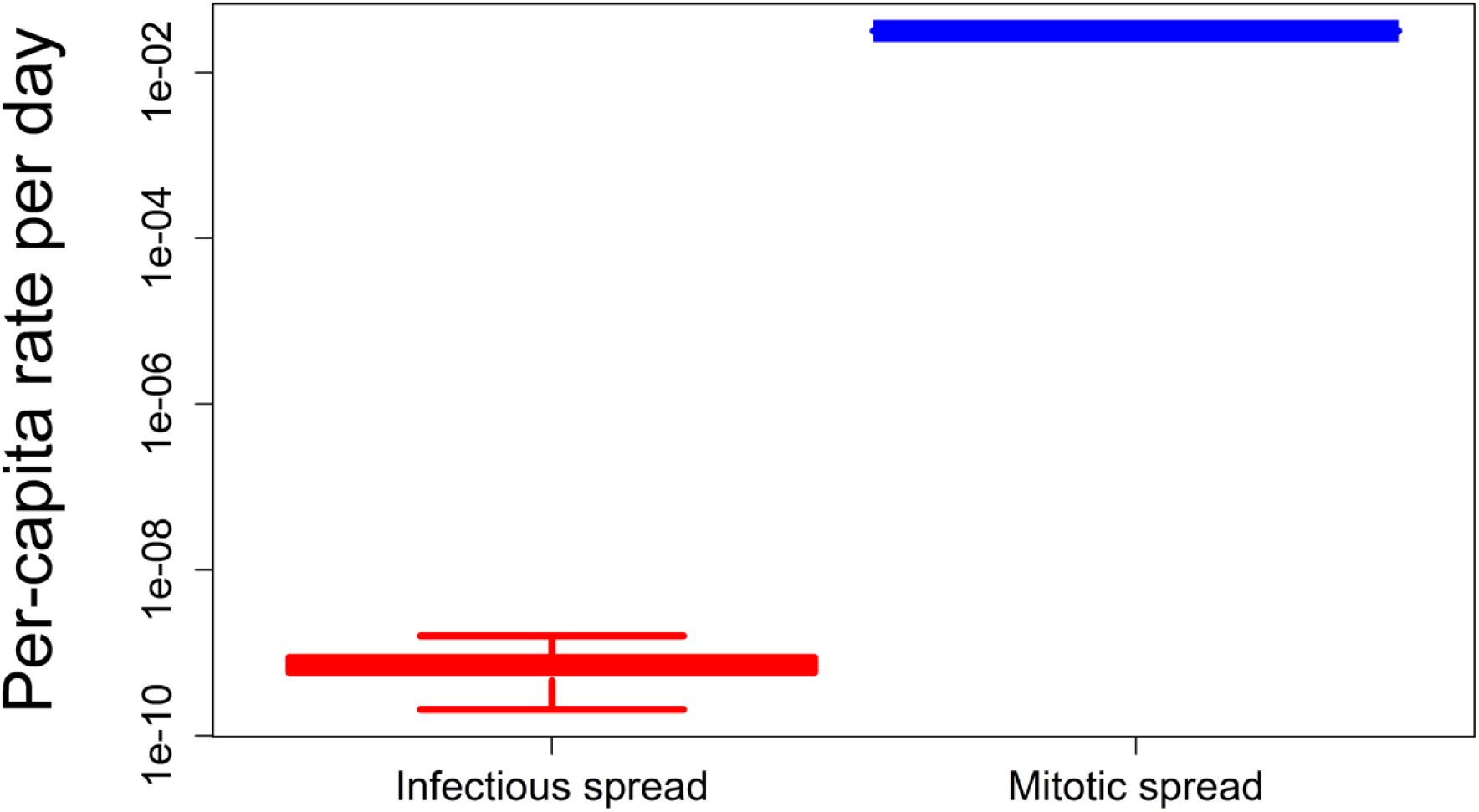
Infectious and mitotic spread rates. Per-capita rates of infectious spread (using hybrid model) and mitotic spread are shown. Infectious spread rates are fitted to HTLV-1 clonal diversity estimates from 11 patients. Mitotic spread rates are derived from previously obtained values [supplementary information]. Mitotic spread is substantially higher than infectious spread in chronic phase of infection.

Within individuals the standard deviation between samples in the infectious spread rate was relatively small, with an average of 2 × 10^−10^ (5.4 × 10^−11^ – 4.1 × 10^−10^) [Table 1]. Estimates of the per-capita infectious spread rate were not found to correlate with either proviral load or with the estimated diversity during the chronic phase (this may be due to our 11 patients providing insufficient power). However, unsurprisingly, the estimated number of new clones per day was correlated with both proviral load (R^2^ = 0.62) and strongly correlated with the estimated diversity (R^2^ = 0.99) [Figure S1].

#### Sensitivity analysis of hybrid model

Originally our threshold value of *F*, above and below which clones are respectively modelled deterministically and stochastically, was set to equal 100. However, the extinction probability of clones of size 100 over a duration of *t*_*Dur*_ = 3133 days [supplementary information] duration was 0.37. We were therefore concerned that excluding such clones would bias the estimates of the infectious spread rate and therefore the ratio, and so re-fitted our model with *F* = 460. This value is the minimum clone frequency for which the extinction probability is less than 1%, given our parameters of infected cell growth, death, and density dependency [Figure S2, supplementary information]. The estimates of infectious spread from the hybrid model are almost identical whether we assume *F* = 100 or *F* = 460. We present the *F* = 460 estimates, as the most accurate description of the system would to consider all clones stochastically. The results of a sensitivity analysis on the length of the time step *h* are shown in Figure S3.

### Modelling approach 2: upper bound approximation

Upper bounds of the infectious spread rate (*r*_*I,Supremum*_) were estimated for each subject using Equation (17), by substituting inputs of HTLV-1 clonal diversity estimates [Table 1] and an estimate of *δ* = 0.0316 infected cell death a day, and an estimate of the total number of infected cells *N* (derived from the proviral load, as detailed in [10]). For each individual we averaged across all samples and across all time points. Estimated values of the rate ranged between individuals from 2.8 × 10^−9^ to 1.7 × 10^−8^ per infected cell per day, and thus (given a rate of per-capita mitotic spread of 0.0316 cells per day) estimates of the ratio *R*_*Supremum*_ ranged between 8.7 × 10^−8^ and 5.5 × 10^−7^ [Figure 6A]. The estimated number of new clones per day using the supremum estimates are unsurprisingly much larger than those of the hybrid, ranging from 516 to 4804, i.e. approximately an order of magnitude higher [Figure 6B].

We further estimated the more restrictive upper bounds of the ratio 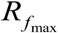 from Equation (21) for multiple *f*_max_ values between 1 and 1000 [Figure 6A]. These estimates assume that the cell death rate applies to clones with frequencies less than or equal to *f*_max_, and that larger clones do not contribute to the rate.

The hybrid estimates always fall below the estimated supremum and are very close to the estimates provided by for *f*_max_ *=* 1 [Figure 6]. Since it is likely that the upper bound approximation will give more accurate estimates for lower values of *f*_max_, this result demonstrates the consistency of estimates produced between the hybrid and the upper bound approximation.

### Modelling approach 3: Occupancy class model

The results from the hybrid model indicate a very low ratio of infectious to mitotic spread. The hybrid benefits from treating small clones stochastically and from the inclusion of known experimental details of HTLV-1 infection and spread. However, it remained possible that these very low estimates of the ratio resulted from incorrect model or parameter assumptions. To test the robustness of our estimate of the ratio to changes in model and parameter assumptions, we adapted a simple deterministic model of HTLV-1 clonal dynamics and occupancy classes and used this to produce two alternative estimators of the ratio of infectious to mitotic spread.

The occupancy class model is based on a model of naïve T cell dynamics developed by de Greef et al [46]. It assumes that clonal dynamics are deterministic, that the clonal structure is in equilibrium and that the probabilities of cell proliferation and death are independent of clone size. The model yields two estimators of the ratio of infectious to mitotic spread. The first estimator (referred to as *R*_*1*_) depends on the proportion of infected cells that are singletons

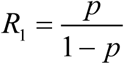

where *p* is the proportion of cells that are singletons.

The second estimator (referred to as *R*_*2*_) depends on species richness.

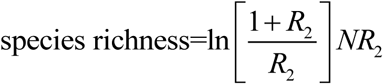

where *N* is the number of infected cells (see Methods for derivation of both expressions).

Across the 99 estimates (11 subjects, 3 time points, 3 replicates) both estimators, *R*_*1*_ and *R*_*2*_, are strongly positively correlated with the estimate of the ratio produced by the hybrid model (P = 1 × 10^−135^ and P = 6 × 10^−87^ respectively, Pearson correlation) and agree well numerically, being of the same order of magnitude and, if anything tending to be even smaller (hybrid median = 2.0 × 10^−8^, hybrid LQ = 1.4 × 10^−8^, hybrid UQ = 3.0 × 10^−8^; *R*_*1*_ median = 2.0 × 10^−8^, *R*_*1*_ LQ=1.4 × 10^−8^, *R*_*1*_ UQ = 3.0 × 10^−8^; *R*_*2*_median = 1.3 × 10^−8^, *R*_*2*_ LQ = 1.0 × 10^−8^, *R*_*2*_ UQ = 1.9 × 10^−8^) [Figure 8].

**Figure 8.**
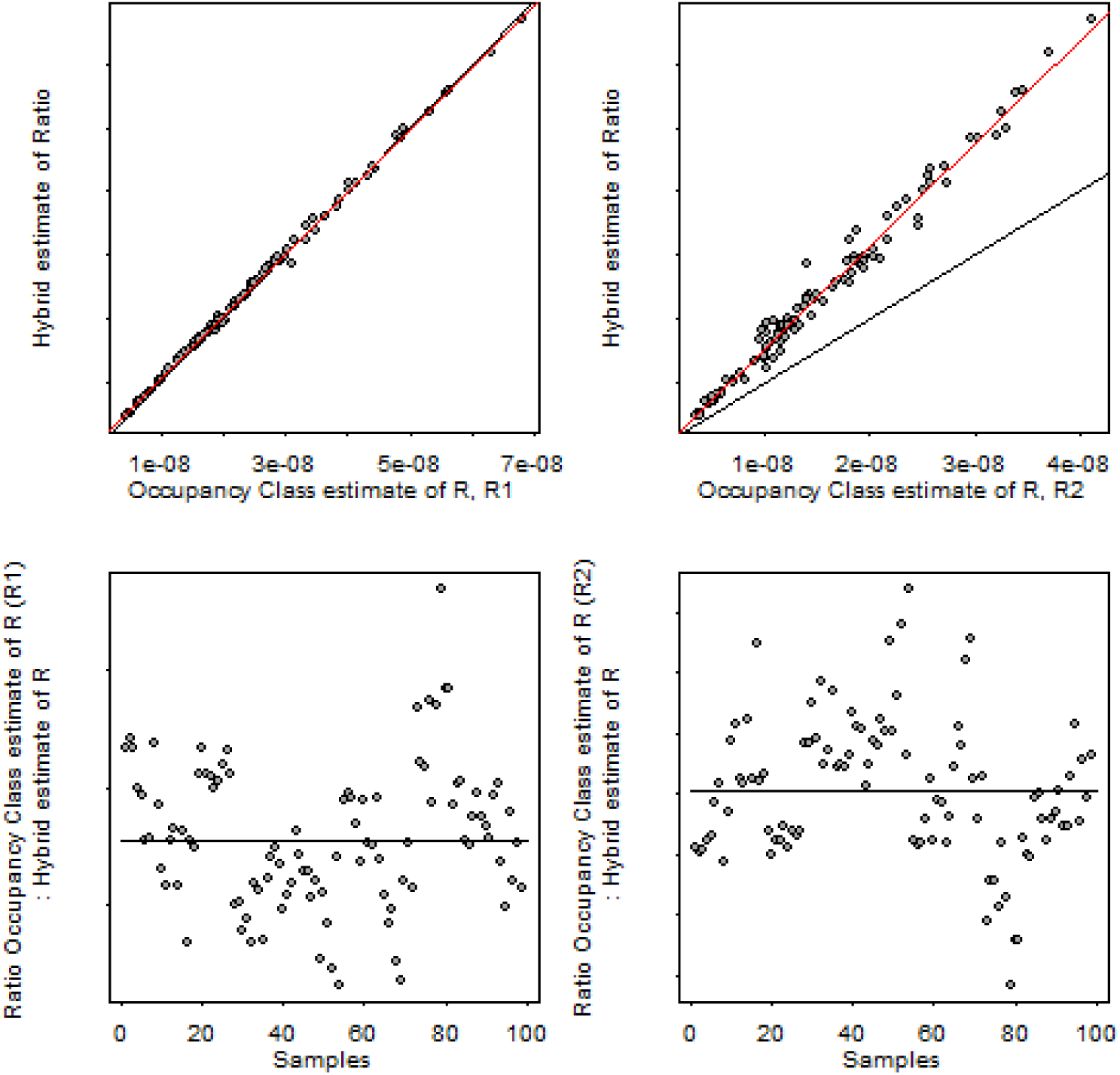
Comparison of estimates of ratio of infectious to mitotic spread from the hybrid model (method 1) and the occupancy class model (method 3). (Top left) Estimate of ratio from hybrid model plotted against first estimate from occupancy class model (*R*_*1*_). Red line is line of best fit, black line is line of equality. (Top right) Estimate of ratio from hybrid model plotted against second estimate from occupancy class model (*R*_*2*_). Red line is line of best fit, black line is line of equality. (Bottom left) Estimate of ratio between hybrid model and first estimate from occupancy class model (*R*_*1*_). Black line denotes the median. (Bottom right) Estimate of ratio between hybrid model and second estimate from occupancy class model (*R*_*2*_). Black line denotes the median.

Finally, we applied the second estimator from the occupancy class model to estimate infectious spread (*R*_*2*_) to the Chao1 estimator of clonal diversity (rather than the DivE estimate used up to this point). The Chao1 estimator gives much lower diversity estimates, and so unsurprisingly yields considerably smaller estimates of the infectious to mitotic spread ratio (median = 7.3 × 10^−10^, LQ = 4.7 × 10^−10^, UQ = 1.0 × 10^−9^).

We conclude that the low estimates of the infectious to mitotic spread are not the product of implicit assumptions in the hybrid model or incorrect parameter choice. Inaccurate estimates of the clonal diversity may play a significant role but calculations using an alternative, widely used estimator provided even smaller estimates of clonal diversity, and therefore yield an even lower ratio.

## Discussion

The relative contribution of infectious and mitotic spread to HTLV-1 viral persistence has not previously been estimated, and this has been a long-standing problem in the field. For many years, it was believed that the virus persisted solely by oligoclonal proliferation of latently infected cells, and that infectious spread contributed little if anything to persistence. However, three observations have brought this belief into question. First, the strong, persistently activated host T-cell response to HTLV-1 implied that the virus is not latent but is frequently expressed in vivo. Second, high-throughput analysis revealed that a typical host carries between 10^4^ and 10^5^ clones, not ∼100 clones as was previously believed. Third, treatment with the antiretroviral therapy lamivudine temporarily but substantially reduced the proviral load of a patient with HAM/TSP. These observations raise the question: what is the contribution of infectious spread to the maintenance of the proviral load during chronic infection?

In this study, we used three different strategies to estimate the ratio of infectious to mitotic spread during the chronic phase of infection. We first developed a deterministic and stochastic hybrid model of within-host HTLV-1 dynamics, and fitted this model to clonal diversity estimates derived from experimental data. We then derived an estimate of the upper bound of the ratio by using a highly simplified model that does not consider individual clones. Finally, we adapted a model of naïve T cell repertoires that models clone occupancy classes. We found broad agreement between the estimates of the ratio obtained using all three methods; and each method implied the existence of ongoing infectious spread during chronic infection, after the HTLV-1 proviral load has reached steady state.

While the ratio of infectious to mitotic spread during the chronic phase is very small (∼2 × 10^−8^), it equates to ∼10^2^ new clones every day. That is, approximately 100 new HTLV-1-infected T cell clones appear every day by infectious spread. Further, while the estimated rate of infectious spread represents a small contribution to overall HTLV-1 persistence, the constant creation of new clones will increase the risk of malignant transformation, because this risk depends in part on the proviral integration site [21]. A malignant clone could originate not only from accumulated mutations in a long-lived clone, but also from a recently infected clone. High HTLV-1 proviral load increases both clonal diversity [47] and risk of ATL [7]. However, it is unknown whether the increased clonal diversity (caused by infectious spread) is a mechanism for this higher risk of malignancy, or whether it is a separate bi-product of high proviral load. Our estimates of ongoing infectious spread during chronic infection are consistent with the hypothesis that higher infectious spread increases the risk of malignant transformation. If this is the case, then anti-retroviral therapy could reduce the risk of ATL in patients who have entered their chronic phase, although it would need to be continued for many years, and would be a long time before its impact was evident.

It is important to note that the different methods we use are not independent. First, they all use our clonal diversity estimates as an input (see section below). Second, they all assume equilibrium clonal diversity. However, they do differ in a number of respects. The upper bound approximation is independent of the parameters *F, π* and *K* and makes no assumptions about the clonal structure or the density dependence of infected cell proliferation. The *R*_*1*_ estimator from the occupancy class model depends only on the proportion of singletons and so is independent of all the parameters (*F, π, δ* and *K*), assumptions about density dependence of proliferation, and indeed the estimated clonal structure beyond the number of singletons. Similarly the *R*_*2*_ estimator from the occupancy class model is also independent of *F, π, δ* and *K* as well as proliferation assumptions. While the hybrid model is our most detailed simulation of HTLV-1 within host dynamics, it is mathematically and computationally complex and requires significant runtime. Because the estimates from all three methods are largely consistent, our analysis indicates that the latter two methods provide good approximations of the rate of infectious spread and the ratio of infectious to mitotic spread.

The most likely source of error in our estimates of the ratio of infectious to mitotic spread lies in the estimation of clonal diversity. Two factors argue against a serious error. First, estimates based on two different quantities (the number of clones and the proportion of infected cells that are singletons) give very similar estimates of the ratio. Second, the DivE estimator compares favourably to other widely-used estimators of species richness [10]. It remains possible that we have underestimated clonal diversity, although it is important to note that DivE produces considerably higher and more plausible estimates than the other estimators, which predicted fewer clones than were observed in additional blood samples taken at the same time.

A much smaller source of potential error lies in using the number of clones to quantify infectious spread. If the virus repeatedly integrates in the same genomic site, then the number of unique genomic sites would be less than the number of true clones, and hence both the infectious spread rate and the ratio would be underestimated. However, hotspots of HTLV-1 integration have not been observed [9], and so such repeat infection would not substantially alter our estimate. Assuming the provirus does not efficiently integrate into heterochromatin, which represents ∼2/3 of the human genome, then only one third of the ∼3 × 10^9^ base pairs of the human genome have the potential for proviral integration. The probability of repeated proviral integration is then the number of existing integration sites divided by the number of potential integration sites. Given the estimated number of clones is of the order of 10^5^, this probability is approximately 10^5^/10^9^ = 10^−4^. Therefore, any error in using the number of clones to quantify infectious spread infectious spread is very small.

It seems surprising that, during initial infection, the virus could establish a stable population of infected T cell clones with such a low rate of infectious spread. However, these low rates of infectious spread are measured in the chronic phase of infection, when the strong host cytotoxic response kills HTLV-1-expressing cells, which probably reduces efficient infectious transmission and favours mitotic transmission. During the early phase of infection, before the establishment of an adaptive immune response, the contribution of infectious spread may be substantially higher than during chronic infection. It would be interesting to model the dynamics of early infection, in particular to investigate the rate required to establish a stable population of infected T cell clones. Modelling early infection would violate the assumption of equilibrium, and thus would void many of the simplifying assumptions that makes our model tractable (e.g. our ability to model clones independently and so avoid an exponential increase in complexity). However, given sufficient computational power, this analysis would be possible.

The methods described here have potential applications in other fields, for example in modelling the human T cell receptor (TCR) repertoire. The mechanisms by which the immune system is reconstituted after immune suppression or transplantation are poorly understood. Drawing parallels between immune reconstitution and HTLV-1 infectious and mitotic spread, the present approach could be applied to investigate the extent to which reconstitution occurs either through the generation of new TCR clonotypes, or through the expansion of existing clonotypes. In HIV-1 infection, the approach could be used to quantify the ratio of infectious to mitotic spread in the absence of treatment and in the latent reservoir remaining following treatment.

In summary, we develop three methods, which have the potential to be applied to a range of areas, and use them to quantify the role of de novo infection in maintaining HTLV-1 viral burden at equilibrium. We find that on average 5 × 10^9^ new infected cells are produced every day; of these the vast majority (>99.9%) will arise from division of an existing infected cell and will thus have the same proviral integration site as their mother cell, but a small minority (about 175 cells per day) will arise from infectious transmission and will contain a novel proviral integration site. These estimates suggest that ongoing infectious spread may be a mechanism for malignant transformation that treatment with antiretroviral drugs may suppress.

## Acknowledgements

We acknowledge grant funding from the Bill and Melinda Gates Foundation, joint Centre funding from the UK Medical Research Council and Department for International Development (grant MR/R015600/1), the Wellcome Trust (Project Grant 091845 to CB, BA; Senior Investigator Award 100291 to CB), and the Imperial College National Institute for Health Research Biomedical Research Centre. We thank the High Performance Computing service staff at Imperial College (https://www.imperial.ac.uk/computational-methods/hpc/), and the staff at the National Centre for Human Retrovirology, Imperial College Healthcare NHS Trust, London, U.K. B.A. is a Wellcome Trust Investigator (103865) and is funded by the Medical Research Council UK (J007439 and G1001052), the European Union Seventh Framework Programme (FP7/2007–2013) under grant agreement 317040 (QuanTI) and Leukemia and Lymphoma Research (15012).

## References

1. Gessain A, Cassar O. Epidemiological Aspects and World Distribution of HTLV-Infection. Frontiers in microbiology. 2012;3:388. Epub 2012/11/20. doi: 10.3389/fmicb.2012.00388. PubMed PMID: 23162541; PubMed Central PMCID: PMC3498738.

2. Ishitsuka K, Tamura K. Human T-cell leukaemia virus type I and adult T-cell leukaemia-lymphoma. Lancet Oncology. 2014;15:e517–e26.

3. Bangham CR, Araujo A, Yamano Y, Taylor GP. HTLV-1-associated myelopathy/tropical spastic paraparesis. Nat Rev Dis Primers. 2015;1:15012. doi: 10.1038/nrdp.2015.12. PubMed PMID: 27188208.

4. Demontis MA, Hilburn S, Taylor GP. Human T cell lymphotropic virus type 1 viral load variability and long-term trends in asymptomatic carriers and in patients with human T cell lymphotropic virus type 1-related diseases. ARHR. 2013;29(2):359–64. Epub 2012/08/17. doi: 10.1089/AID.2012.0132. PubMed PMID: 22894552.

5. Matsuzaki T, Nakagawa M, Nagai M, Usuku K, Higuchi I, Arimura K, et al. HTLV-I proviral load correlates with progression of motor disability in HAM/TSP: analysis of 239 HAM/TSP patients including 64 patients followed up for 10 years. Journal of neurovirology. 2001;7(3):228-34. PubMed PMID: 11517397.

6. Nagai M, Usuku K, Matsumoto W, Kodama D, Takenouchi N, Moritoyo T, et al. Analysis of HTLV-I proviral load in 202 HAM/TSP patients and 243 asymptomatic HTLV-I carriers: high proviral load strongly predisposes to HAM/TSP. J Neurovirol. 1998;4(6):586–93. Epub 1999/03/05. PubMed PMID: 10065900.

7. Okayama A, Stuver S, Matsuoka M, Ishizaki J, Tanaka G, Kubuki Y, et al. Role of HTLV-1 proviral DNA load and clonality in the development of adult T-cell leukemia/lymphoma in asymptomatic carriers. Int J Cancer. 2004;110(4):621–5. Epub 2004/05/04. doi: 10.1002/ijc.20144. PubMed PMID: 15122598.

8. Overbaugh J, Bangham CR. Selection forces and constraints on retroviral sequence variation. Science. 2001;292(5519):1106–9. Epub 2001/05/16. PubMed PMID: 11352065.

9. Gillet NA, Malani N, Melamed A, Gormley N, Carter R, Bentley D, et al. The host genomic environment of the provirus determines the abundance of HTLV-1-infected T-cell clones. Blood. 2011;117(11):3113–22. Epub 2011/01/14. doi: blood-2010-10-312926 [pii] 10.1182/blood-2010-10-312926. PubMed PMID: 21228324; PubMed Central PMCID: PMC3062313.

10. Laydon DJ, Melamed A, Sim A, Gillet NA, Sim K, Darko S, et al. Quantification of HTLV-1 clonality and TCR diversity. PLoS computational biology. 2014;10(6):e1003646. doi: 10.1371/journal.pcbi.1003646. PubMed PMID: 24945836; PubMed Central PMCID: PMC4063693.

11. Wattel E, Vartanian JP, Pannetier C, Wain-Hobson S. Clonal expansion of human T-cell leukemia virus type I-infected cells in asymptomatic and symptomatic carriers without malignancy. J Virol. 1995;69(5):2863–8. Epub 1995/05/01. PubMed PMID: 7707509; PubMed Central PMCID: PMC188982.

12. Tanaka G, Okayama A, Watanabe T, Aizawa S, Stuver S, Mueller N, et al. The clonal expansion of human T lymphotropic virus type 1-infected T cells: a comparison between seroconverters and long-term carriers. J Infect Dis. 2005;191(7):1140–7. Epub 2005/03/05. doi: JID33135 [pii] 10.1086/428625. PubMed PMID: 15747250.

13. Wattel E, Cavrois M, Gessain A, Wain-Hobson S. Clonal expansion of infected cells: a way of life for HTLV-I. J Acquir Immune Defic Syndr Hum Retrovirol. 1996;13 Suppl 1:S92–9. Epub 1996/01/01. PubMed PMID: 8797710.

14. Wodarz D, Nowak MA, Bangham CR. The dynamics of HTLV-I and the CTL response. Immunol Today. 1999;20(5):220–7. Epub 1999/05/14. doi: S0167569999014462 [pii]. PubMed PMID: 10322301.

15. Berry CC, Gillet NA, Melamed A, Gormley N, Bangham CR, Bushman FD. Estimating abundances of retroviral insertion sites from DNA fragment length data. Bioinformatics. 2012;28(6):755–62. Epub 2012/01/13. doi: bts004 [pii] 10.1093/bioinformatics/bts004. PubMed PMID: 22238265; PubMed Central PMCID: PMC3307109.

16. Cavrois M, Wain-Hobson S, Gessain A, Plumelle Y, Wattel E. Adult T-cell leukemia/lymphoma on a background of clonally expanding human T-cell leukemia virus type-1-positive cells. Blood. 1996;88(12):4646–50. Epub 1996/12/15. PubMed PMID: 8977257.

17. Furukawa Y, Fujisawa J, Osame M, Toita M, Sonoda S, Kubota R, et al. Frequent clonal proliferation of human T-cell leukemia virus type 1 (HTLV-1)-infected T cells in HTLV-1-associated myelopathy (HAM-TSP). Blood. 1992;80(4):1012–6. Epub 1992/08/15. PubMed PMID: 1498321.

18. Gabet AS, Mortreux F, Talarmin A, Plumelle Y, Leclercq I, Leroy A, et al. High circulating proviral load with oligoclonal expansion of HTLV-1 bearing T cells in HTLV-1 carriers with strongyloidiasis. Oncogene. 2000;19(43):4954–60. Epub 2000/10/24. doi: 10.1038/sj.onc.1203870. PubMed PMID: 11042682.

19. Meekings KN, Leipzig J, Bushman FD, Taylor GP, Bangham CR. HTLV-1 integration into transcriptionally active genomic regions is associated with proviral expression and with HAM/TSP. PLoS Pathog. 2008;4(3):e1000027. Epub 2008/03/29. doi: 10.1371/journal.ppat.1000027. PubMed PMID: 18369476; PubMed Central PMCID: PMC2265437.

20. Bangham CR. Human T-cell leukaemia virus type I and neurological disease. Curr Opin Neurobiol. 1993;3(5):773–8. Epub 1993/10/01. PubMed PMID: 8260828.

21. Cook LB, Melamed A, Niederer H, Valganon M, Laydon D, Foroni L, et al. The role of HTLV-1 clonality, proviral structure, and genomic integration site in adult T-cell leukemia/lymphoma. Blood. 2014;123(25):3925–31. doi: 10.1182/blood-2014-02-553602. PubMed PMID: 24735963; PubMed Central PMCID: PMC4064332.

22. Gillet NA, Cook L, Laydon DJ, Hlela C, Verdonck K, Alvarez C, et al. Strongyloidiasis and infective dermatitis alter human T lymphotropic virus-1 clonality in vivo. PLoS Path. 2013;9(4):e1003263. Epub 2013/04/18. doi: 10.1371/journal.ppat.1003263 PPATHOGENS-D-12-02814 [pii]. PubMed PMID: 23592987; PubMed Central PMCID: PMC3617147.

23. Taylor GP, Hall SE, Navarrete S, Michie CA, Davis R, Witkover AD, et al. Effect of lamivudine on human T-cell leukemia virus type 1 (HTLV-1) DNA copy number, T-cell phenotype, and anti-tax cytotoxic T-cell frequency in patients with HTLV-1-associated myelopathy. Journal of virology. 1999;73(12):10289–95.

24. Laydon DJ, Bangham CR, Asquith B. Estimating T-cell repertoire diversity: limitations of classical estimators and a new approach. Phil Trans R Soc Lond B. 2015;370(1675). doi: 10.1098/rstb.2014.0291. PubMed PMID: 26150657; PubMed Central PMCID: PMCPMC4528489.

25. Laydon DJ, Sim A, Bangham CRM, Asquith B. DivE: Diversity Estimator. 1.1 ed2019.

26. Jahnke T, Kreim M. Error bound for piecewise deterministic processes modeling stochastic reaction systems. Multiscale Modeling & Simulation. 2012;10(4):1119–47.

27. Jahnke T, Sunkara V. Error Bound for Hybrid Models of Two-Scaled Stochastic Reaction Systems. In: Dahlke S, Dahmen W, Griebel M, Hackbusch W, Ritter K, Schneider R, et al., editors. Extraction of Quantifiable Information from Complex Systems. Cham: Springer International Publishing; 2014. p. 303–19.

28. Gillespie DT. A rigorous derivation of the chemical master equation. Physica A: Statistical Mechanics and its Applications. 1992;188(1):404–25.

29. Jahnke T. On reduced models for the chemical master equation. Multiscale Modeling & Simulation. 2011;9(4):1646–76.

30. Van Kampen NG. Stochastic processes in physics and chemistry: Elsevier; 1992.

31. Hegland M, Burden C, Santoso L, MacNamara S, Booth H. A solver for the stochastic master equation applied to gene regulatory networks. Journal of computational and applied mathematics. 2007;205(2):708–24.

32. Jahnke T, Huisinga W. A dynamical low-rank approach to the chemical master equation. Bulletin of mathematical biology. 2008;70(8):2283–302.

33. Stewart WJ. Introduction to the numerical solutions of Markov chains: Princeton Univ. Press; 1994.

34. Strang G. On the construction and comparison of difference schemes. SIAM Journal on Numerical Analysis. 1968;5(3):506–17.

35. Asquith B, McLean AR. In vivo CD8+ T cell control of immunodeficiency virus infection in humans and macaques. PNAS. 2007;104(15):6365–70. Epub 2007/04/04. doi: 0700666104 [pii] 10.1073/pnas.0700666104. PubMed PMID: 17404226.

36. Chao A. Nonparametric estimation of the number of classes in a population. Scandinavian Journal of Statistics. 1984;11(4):265–70.

37. La Gruta NL, Rothwell WT, Cukalac T, Swan NG, Valkenburg SA, Kedzierska K, et al. Primary CTL response magnitude in mice is determined by the extent of naive T cell recruitment and subsequent clonal expansion. The Journal of Clinical Investigation. 2010;120(6):1885–94.

38. Gwinn DC, Allen MS, Bonvechio KI, V. Hoyer M, Beesley LS. Evaluating estimators of species richness: the importance of considering statistical error rates. Methods in Ecology and Evolution. 2016;7(3):294–302.

39. Hamad I, Ranque S, Azhar EI, Yasir M, Jiman-Fatani AA, Tissot-Dupont H, et al. Culturomics and amplicon-based metagenomic approaches for the study of fungal population in human gut microbiota. Scientific reports. 2017;7(1):16788.

40. Branco M, Figueiras FG, Cermeño P. Assessing the efficiency of non-parametric estimators of species richness for marine microplankton. Journal of Plankton Research. 2018;40(3):230–43.

41. R Core Team. R: A language and environment for statistical computing [Internet]. Vienna, Austria; 2018. 3.5.0 ed. Vienna, Austria: R Foundation for Statistical Computing; 2018.

42. Dowle M, Srinivasan A, Gorecki J, Short T, Lianoglou S, Antonyan E. data. table: extension of data. frame. R package version 1.9. 8. 2016. 2017.

43. Bates D, Maechler M, Maechler MM. Package ‘Matrix’. 2017.

44. Arioli M, Codenotti B, Fassino C. The Padé method for computing the matrix exponential. Linear algebra and its applications. 1996;240:111–30.

45. Brent RP. Algorithms for minimization without derivatives: Courier Corporation; 2013.

46. de Greef PC, Oakes T, Gerritsen B, Ismail M, Heather JM, Hermsen R, et al. The naive T-cell receptor repertoire has an extremely broad distribution of clone sizes. bioRxiv. 2019:691501. doi: 10.1101/691501.

47. Niederer HA, Laydon DJ, Melamed A, Elemans M, Asquith B, Matsuoka M, et al. HTLV-1 proviral integration sites differ between asymptomatic carriers and patients with HAM/TSP. Virology journal. 2014;11(1):172.

48. Asquith B, Zhang Y, Mosley AJ, de Lara CM, Wallace DL, Worth A, et al. In vivo T lymphocyte dynamics in humans and the impact of human T-lymphotropic virus 1 infection. Proc Natl Acad Sci U S A. 2007;104(19):8035–40. Epub 2007/05/08. doi: 0608832104 [pii] 10.1073/pnas.0608832104. PubMed PMID: 17483473; PubMed Central PMCID: PMC1861853.

